# A Single-Molecule Long-Read Survey of Human Transcriptomes using LoopSeq Synthetic Long Read Sequencing

**DOI:** 10.1101/532135

**Authors:** Indira Wu, Hee Shin Kim, Tuval Ben-Yehezkel

**Affiliations:** Loop Genomics, San Jose, CA, 95138

**Author notes:** Corresponding author: Tuval Ben-Yehezkel.

## Abstract

State-of-the-art short-read transcriptome sequencing methods employ unique molecular identifier (UMI) to accurately classify and count mRNA transcripts. A fundamental limitation of UMI-based short-read transcriptome sequencing is that each read typically covers a small fraction of the transcript sequence. Efforts to accurately characterize splicing isoforms, arguably the largest source of variation in Human gene expression, using short read sequencing have therefore largely relied on computational predictions of transcript isoforms based on indirect observations. Here we describe a transcript counting, synthetic long read method for sequencing whole transcriptomes using short read sequencing platforms and no additional hardware. The method enables full-length mRNA sequence reconstruction at single-nucleotide resolutions with high-throughput, low error rates and UMI based transcript counting using any Illumina sequencer. We describe results from whole transcriptome sequencing from total RNA extracted from 3 human tissue samples: brain, liver, and blood. Reconstructed transcript sequences are characterized and annotated using SQANTI, an analysis pipeline for assessing the sequence quality of long-read transcriptomes. Our results demonstrate that LoopSeq synthetic long-read sequencing can reconstruct contigs up to 3,900nt full-length transcripts using tissue extracted RNA, as well as identify novel splice variants of known junction donors and acceptors.

## Introduction

Whole transcriptome analysis has a profound impact in understanding the relationship between gene expressions, isoform sequences, and the complex phenotypes of cellular developments and diseases. With the advance in Next Generation Sequencing, many different protocols and toolsets have been developed to provide more insights into whole transcriptome at the sequence level. Current transcriptome sequencing techniques include 3’ short read sequencing of polyadenylated RNA while measuring transcript abundance, reference-based short read transcript sequence assembly, and full-length transcript sequencing using long-read sequencers. While current transcriptome sequencing methods have made large strides in the transcriptome field, it remains difficult and cost-prohibitive to detect and discover new alternative splicing events, which have been implicated in many cancers and hereditary diseases.

We have developed a method for reconstructing long read sequences using Illumina’s short-read sequencers. The sample prep workflow includes reverse transcription, cDNA amplification, barcode distribution, and Illumina adaptor attachment and amplification. During reverse transcription, each complementary DNA (cDNA) is tagged with sample barcode and a unique molecular identifier (UMI). The UMI, or barcode, is used to differentiate between PCR duplicates and transcripts of identical sequences, as well as providing relative abundance of the transcripts without the interference of PCR amplification bias. Following cDNA amplification, the barcodes on each cDNA is distributed across the length of the cDNA molecule, before being converted into Illumina sequencing libraries complete with handles suitable for cluster amplification and sequencing on an Illumina sequencer. With a simple modification of the sample prep workflow, the short-read sequencing read coverage can either be used for reconstructing long RNA transcripts (“phasing” workflow) or for counting transcripts with a sparse coverage within each transcript (“counting” workflow). The choice of workflow offers flexibility for when transcript discovery and isoform identification are the focus of the study or when transcriptome abundance measurement is desired. In the data presented below, the transcriptomes of total RNA extracted from human brain, liver, and blood are surveyed at different sequencing depth per transcript. The impact of RNA sample integrity as well as sequencing depth on the reconstructed long reads are examined. Sequence annotation and splice junction analysis are conducted using SQANTI [1], an analysis pipeline for characterizing long read transcript sequences.

## Results

### Input RNA Analysis

The RNA integrity of the total RNA from human brain, liver, and blood were assessed using Agilent Bioanalyzer RNA 6000 Nano kit coupled with the Eukaryote Total RNA analysis workflow. Figure 1 shows the electropherograms of different human total RNA samples. The highest RIN score of the total RNA sample is from human liver total RNA at 7.4, followed by the human brain total RNA with a RIN score of 6.1, followed by the human blood total RNA with a RIN score of 2.4. The three human total RNA samples cover a wide range of RIN scores at realistic sample integrities that one might encounter with tissue extracted RNA and serve as good model samples for method validation with the LoopSeq Transcriptome kit.

**Figure 1.**
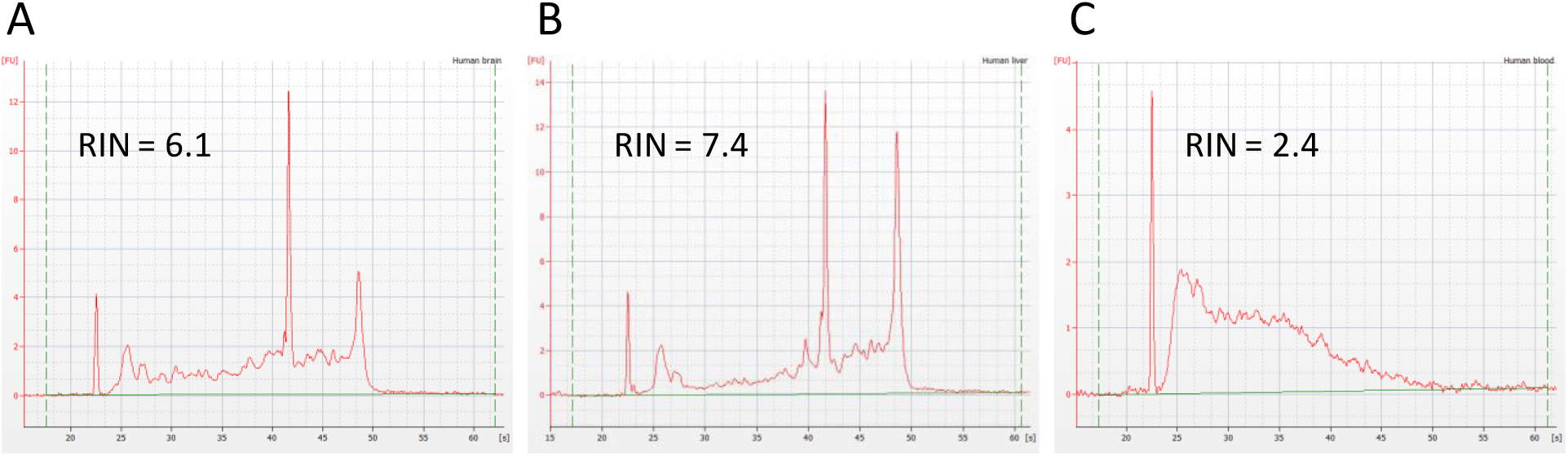
Electrophoretic separation of different human total RNA using the RNA 6000 Nano Kit with the Eukaryote Total RNA assay. The position of 18S and 25S rRNA peaks are marked, and the RIN scores are shown. Unlabeled peaks correspond to ribosomal RNA degradation products. A) Human brain total RNA. B) Human liver total RNA. C) Human blood total RNA.

### Relationship between Sample Complexity and Sequencing Depth

For *de* novo assembly of RNA transcripts, it is important to consider the sequencing fold coverage one needs to sufficiently cover the starting molecule, or “coverage per transcript” as referred to below. When considering the sequencing need on a per sample level, one also needs to consider the number of molecules the sequencing read coverage is going towards. Different total RNA samples can contain different levels of mRNA molecules, which in terms impacts the number of uniquely barcoded cDNA molecules that are made. This is referred to as the “sample complexity” of the tagged sample, or the number of uniquely barcoded molecules that can be sequenced. The same number of paired-end short reads can be used towards sequencing different sample prep libraries with different sample complexities. An example of the impact on contig lengths from two different sequencing coverages per transcript is shown in Figure 2. Two sequencing libraries for the human brain RNA sample were constructed, using different numbers of uniquely tagged cDNA as inputs into the cDNA amplification reactions. For the medium coverage per transcript library, 6.7M PE150 reads were used to reconstruct ∼23,000 unique transcripts, while for the high coverage per transcript library, 5.2M PE150 reads were used reconstruct ∼5,200 unique transcripts. The average contig length from a medium coverage per transcript is ∼700bp, in contrast with the average contig length of ∼1300bp from a high sequencing coverage per transcript. Additionally, the high coverage per transcript dataset include contigs up to 3900bp, and roughly 30% of the transcripts cover >80% of the reference sequences. The contig sequences and the contig mapping statistics for the human brain sample can be downloaded <u>here</u> for the high coverage per transcript data and <u>here</u> for the medium coverage per transcript data.

**Figure 2.**
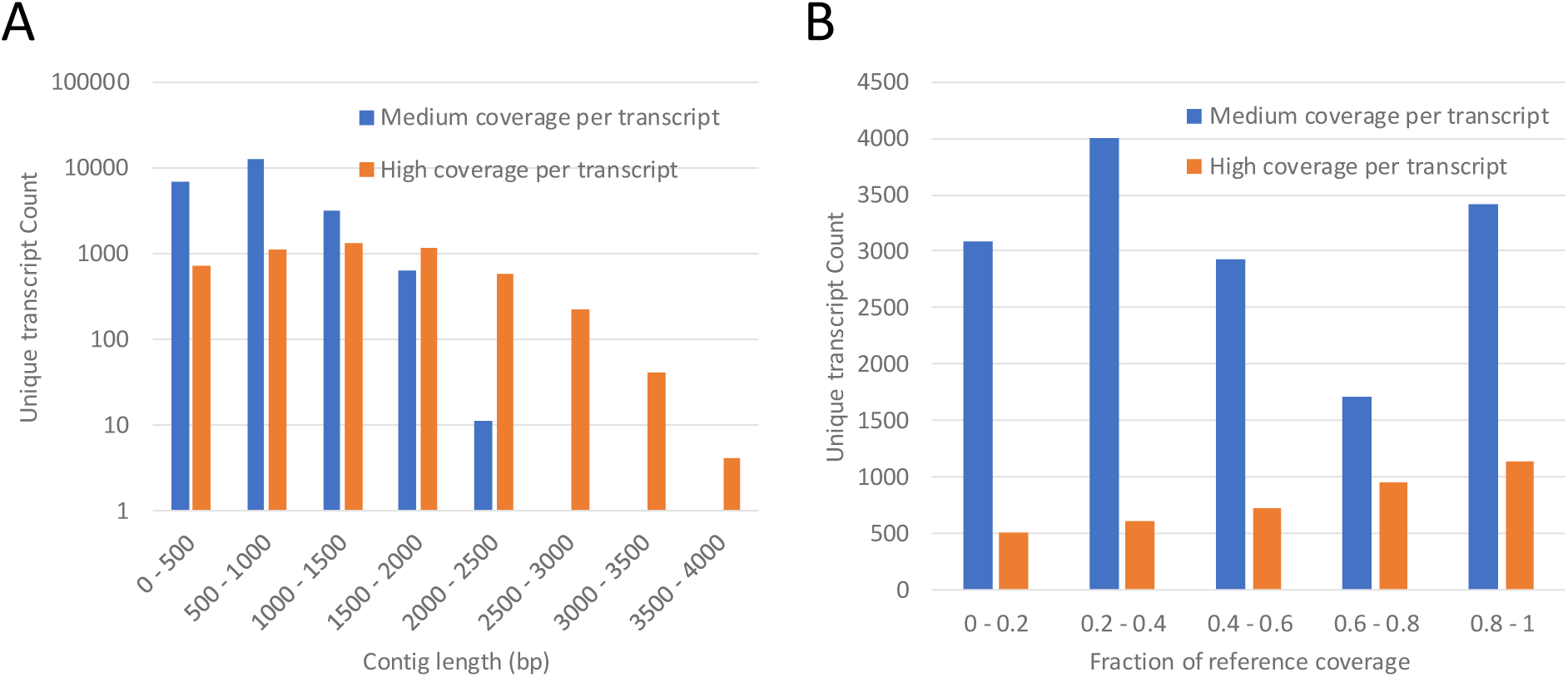
The effect of sequencing depth on the reconstructed transcript lengths. A) The contig length histograms of unique transcripts from two different sequencing coverages of the human brain sample. B) The fraction of reference coverage from two different sequence coverages of the human brain sample.

At the extreme ends, when the short-read coverage is 20 reads or lower per transcript, *e.g.* with the counting workflow of the LoopSeq Transcriptome kit, the dataset is best used for counting transcripts. With the sparse coverage per transcript, one can still obtain information on the gene functions of the transcripts but not necessarily the transcript sequence in its entirely. On the other hand, when the short-read coverage is 200 reads or more per transcript, *e.g.* with the phasing workflow of the LoopSeq Transcriptome kit, the dataset may be used to reconstruct transcript sequences. Depending on the length of the starting RNA molecules, that can either lead to a full-length reconstruction with short RNA molecules or a partial reconstruction with long RNA molecules. Additional short read coverage contributes more sequence information disproportionally towards longer starting RNA molecules, leading to longer contig reconstruction and full length contigs.

### Impact of Sample Integrity and Contig Length

During reverse transcription and molecular barcode tagging, the RNA molecules are copied into complementary DNA (cDNA) for downstream sample preps. Only full-length RNA molecules can be reverse transcribed into cDNA and amplified. When the starting RNA sample is highly degraded, the length distribution of the reconstructed contigs are generally short because the cDNAs that were tagged and amplified were short. An example of the impact on contig length due to the RNA sample integrity is shown in Figure 3. Synthetic long read sequencing libraries for the human brain, liver, and blood RNA samples were constructed, using roughly 20,000 to 30,000 uniquely tagged cDNA as inputs into the cDNA amplification reactions. Between the human brain sample (RIN score of 6.1) and the human liver sample (RIN score of 7.4), the contig length histogram as well as the fraction of reference coverage are comparable. However, for the human blood sample, with a RIN score of 2.1 indicating significant RNA degradation, the reconstructed contigs are mostly 1000bp or shorter, in contrast with the human brain and the human liver samples which have contigs up to 2500bp. When examining the reference coverage by the reconstructed contigs, there is a roughly equal distribution across all possible reference coverage in length for the human brain and the human liver sample, while more than 50% of the contigs cover 80% of the references in length with the human blood sample. With majority of the contigs covering close to full length transcripts, this indicates that the contig lengths of the human blood sample, though mostly 1000bp or shorter, represent the true transcript lengths in the starting RNA sample. Since only full-length RNA molecules can be prepared into sequencing library with the LoopSeq Transcriptome kit, the RNA sample degradation directly translates to the absence of long transcript sequences. The contig sequences and the contig mapping statistics for the human brain, liver, and blood samples with medium coverage per transcript can be downloaded <u>here</u>, <u>here</u>, and <u>here</u>, respectively.

**Figure 3.**
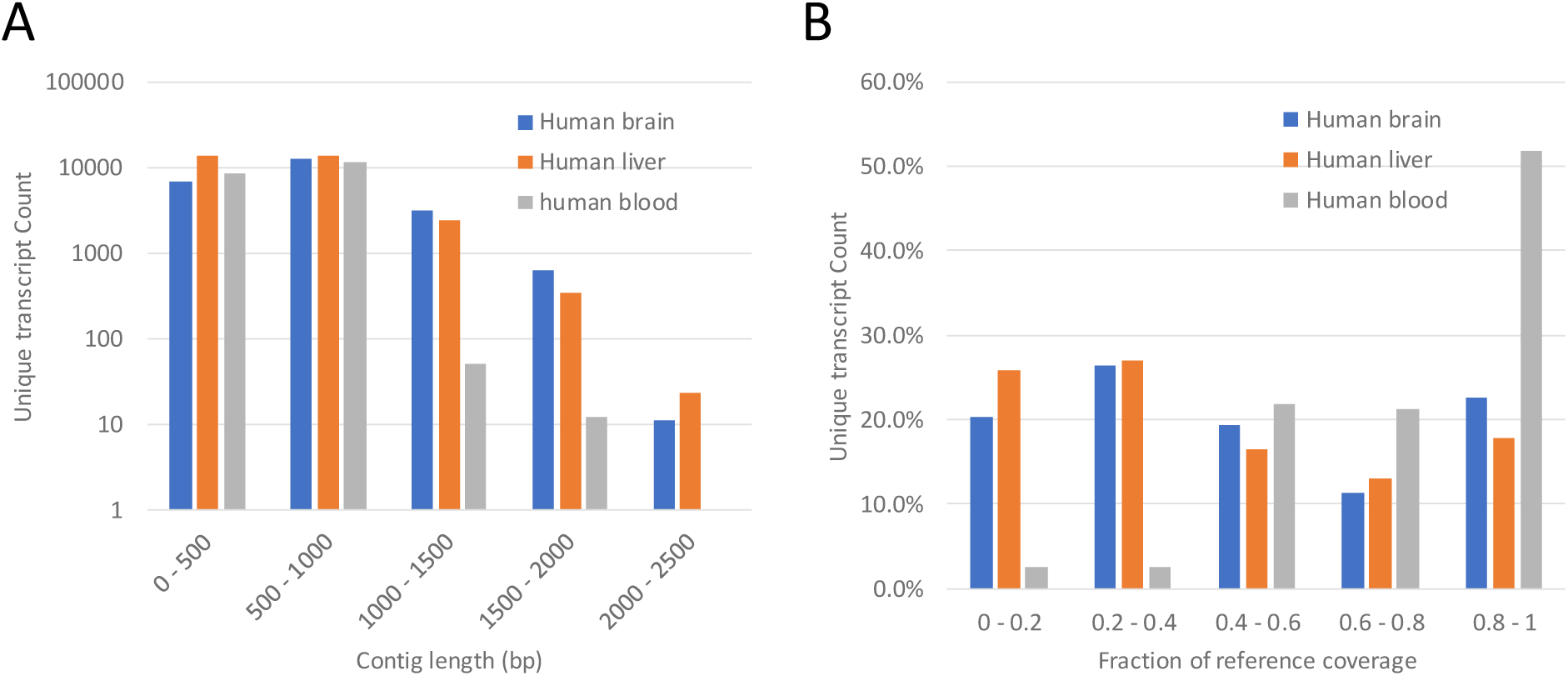
The impact of sample integrity on the contig lengths of different human RNA samples at medium coverage per transcript. A) The contig length histogram from 3 different human RNA samples: brain, liver, and blood. B) The fraction of reference coverage from the 3 different human RNA samples: brain, liver, and blood. A 0.8 fraction of reference coverage means of the reference that each contig selects as the best match for its sequence, 80% of the said reference length is covered by the contig.

### Long-read Transcript Analysis using SQANTI

To analyze the single-molecule long reads, the contigs of the human brain and the human liver samples were taken through the SQANTI pipeline for transcript quality assessment and transcript classification. The SQANTI report and output files can be downloaded <u>here</u> for the human brain sample and <u>here</u> for the human liver. By comparing the splice junctions observed in the contig sequences against the human reference genome GRCh38, the quality of the reconstructed contigs were assessed. The annotated contigs, or isoforms, are summarized in Table 1. SQANTI analysis annotates the contigs using the following criteria:

**Table 1.**
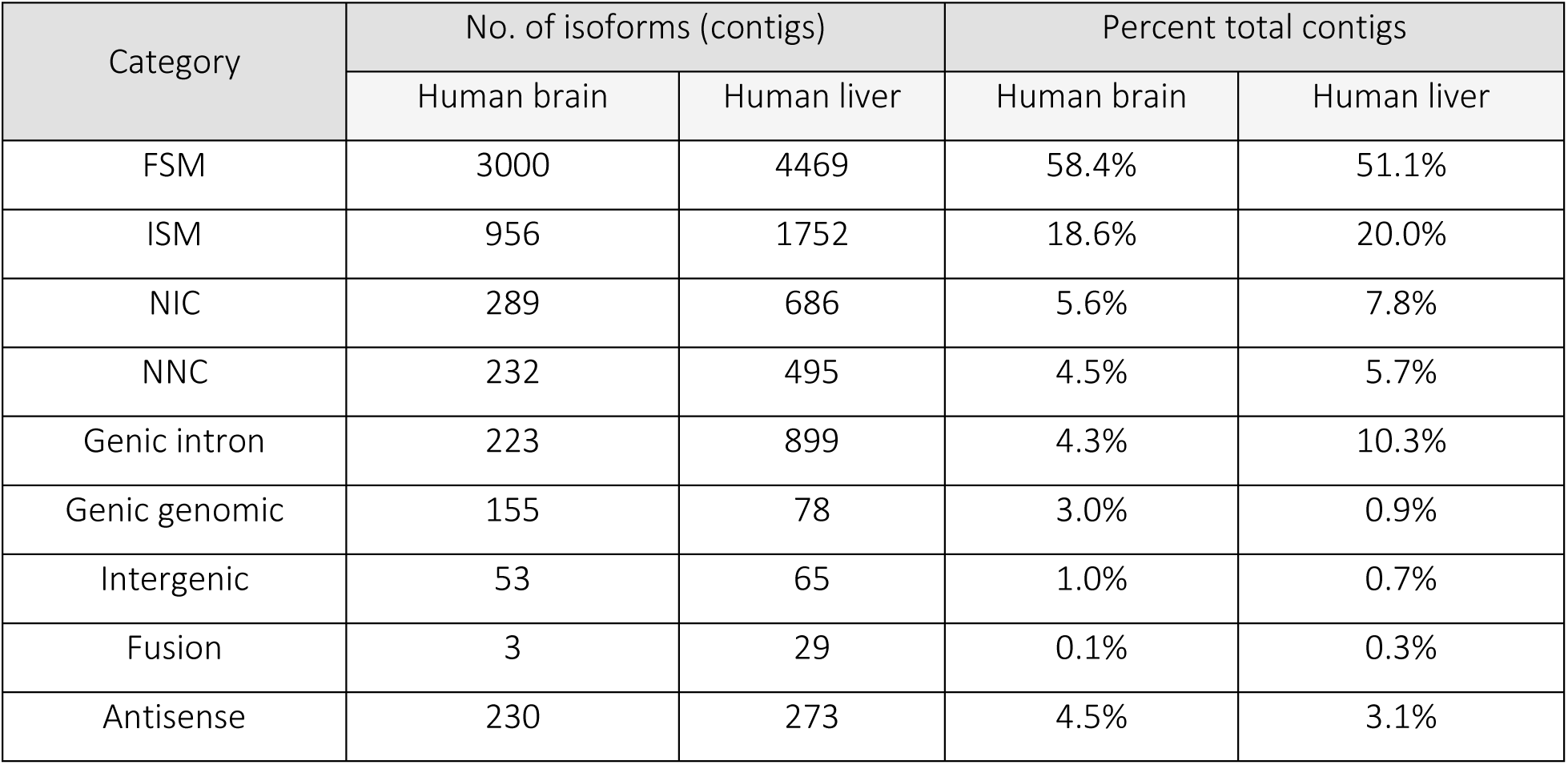
SQUANTI annotations of the contigs from different human RNA samples.

- Full Splice Match (FSM): the input contigs match the reference transcripts at all splice junctions
- Incomplete Splice Match (ISM): the input contigs match consecutive but not all splice junctions of the reference transcripts
- Novel in Category (NIC): the input contigs contain new combinations of previously annotated splice junctions, or novel splice junctions from already annotated junction donors and acceptors
- Novel not in Category (NNC): the input contigs contain novel junction donor and/or acceptors of previously annotated genes
- Intergenic: the input contigs lies outside the boundaries of an annotated gene
- Genic intron: the input contigs lies entirely within the boundaries of an annotated intron
- Genic genomic: the input contigs include partial exon and intron/intergenic region of an annotated gene
- Fusion: the input contigs spans two annotated loci
- Antisense: the input contigs contain sequences that are the complementary strand of an annotated transcript

Using the SQANTI annotation of the contigs, isoforms mapping to a known reference (FSM, ISM) account for 77.0% and 71.1% of the reconstructed contigs for human brain and liver RNA, respectively, while novel isoforms of known genes account for another 10.1% and 13.5% of the reconstructed contigs. Genic introns, which are contigs that map entirely within the boundaries of an annotated intron, account for the majority of the remaining reconstructed contigs. Antisense RNA, where contigs map to the reverse complement sequence of an annotated gene, is another important type of transcript isoform. Since LoopSeq assigns the UMI during reverse transcription when the first strand cDNA is synthesized, the LoopSeq reconstructed long read is strand-aware, or stranded. For the human brain and liver samples, SQANTI reports 3-5% of the reconstructed contigs to be of the antisense strand. Lastly, the LoopSeq reconstructed contigs may or may not be full length transcripts because contig lengths are highly dependent on the sequencing depth of the sample and the sequence coverage per transcript, as discussed previously (Figure 2).

To dissect the novel isoforms further, SQANTI analysis also examines the splice junctions of the contigs. Two categories of splice junctions are defined using the dinucleotides at the beginning and the end of the introns that are spliced:

- Canonical: GT-AG, GC-AG, AT-AC dinucleotide pairs. This arises from the observation that 99.9% of all human introns are composed of these 3 dinucleotide pairs, with GT-AG being the most common dinucleotide pair in the human genome.
- Non-canonical: all other possible combination not include as canonical dinucleotide pairs.

Examining the splice junctions in the reconstructed contigs, canonical splice junctions from previously annotated genes account for 92.8% and 89.9% of the observed splice junctions with the human brain and human liver sample respectively, while non-canonical splice junctions from previously annotated genes account for less than 0.1% of the splice junctions. Novel canonical splice junctions, representing new combination of junction donors and acceptors with canonical splice junctions, account for 2.2% and 3.8% of the splice junctions with human brain and human liver sample. Novel non-canonical splice junctions, representing new junction donors and acceptors with non-canonical splice junctions, account for 4.9% and 6.2% of the splice junctions for human brain and human liver sample. Note that non-canonical splice junctions are more prevalent with novel genes than with previously annotated genes. Given that canonical splice junctions account for 99.9% of all human introns, it is believed that one source of the non-canonical splice junctions is due to the template switching property of the reverse transcriptase [2, 3]. To detect reverse transcriptase switching event, SQANTI implements an algorithm searching for the presence of repeat sequence between the upstream boundary of the non-canonical intron and the intron region adjacent to the downstream exon boundary. The presence of such repeat sequence would indicate that template switching by the reverse transcriptase may take place, resulting in non-canonical splice junctions. As shown in Figure 4, the RT switching event categorized by the presence of repeat sequence is found at roughly 7 – 10% of the splice junctions in novel transcripts (canonical and non-canonical), while it is found <1% of the splice junctions in known transcripts. The higher prevalence of repeat sequence in novel transcripts suggest up to 10% of the novel transcripts may arise from an artifact of the reverse transcriptase during sample preparation.

**Table 2.** SQUANTI splice junction analysis of the contigs from different human RNA samples.

| Category | No. of splice junctions |  | Percent total splice junctions |  |
| --- | --- | --- | --- | --- |
|  | Human brain | Human liver | Human brain | Human liver |
| Known canonical | 8702 | 10006 | 92.8% | 89.9% |
| Known non-canonical | 6 | 3 | 0.1% | 0.0% |
| Novel canonical | 205 | 421 | 2.2% | 3.8% |
| Novel non-canonical | 462 | 694 | 4.9% | 6.2% |

**Figure 4.**
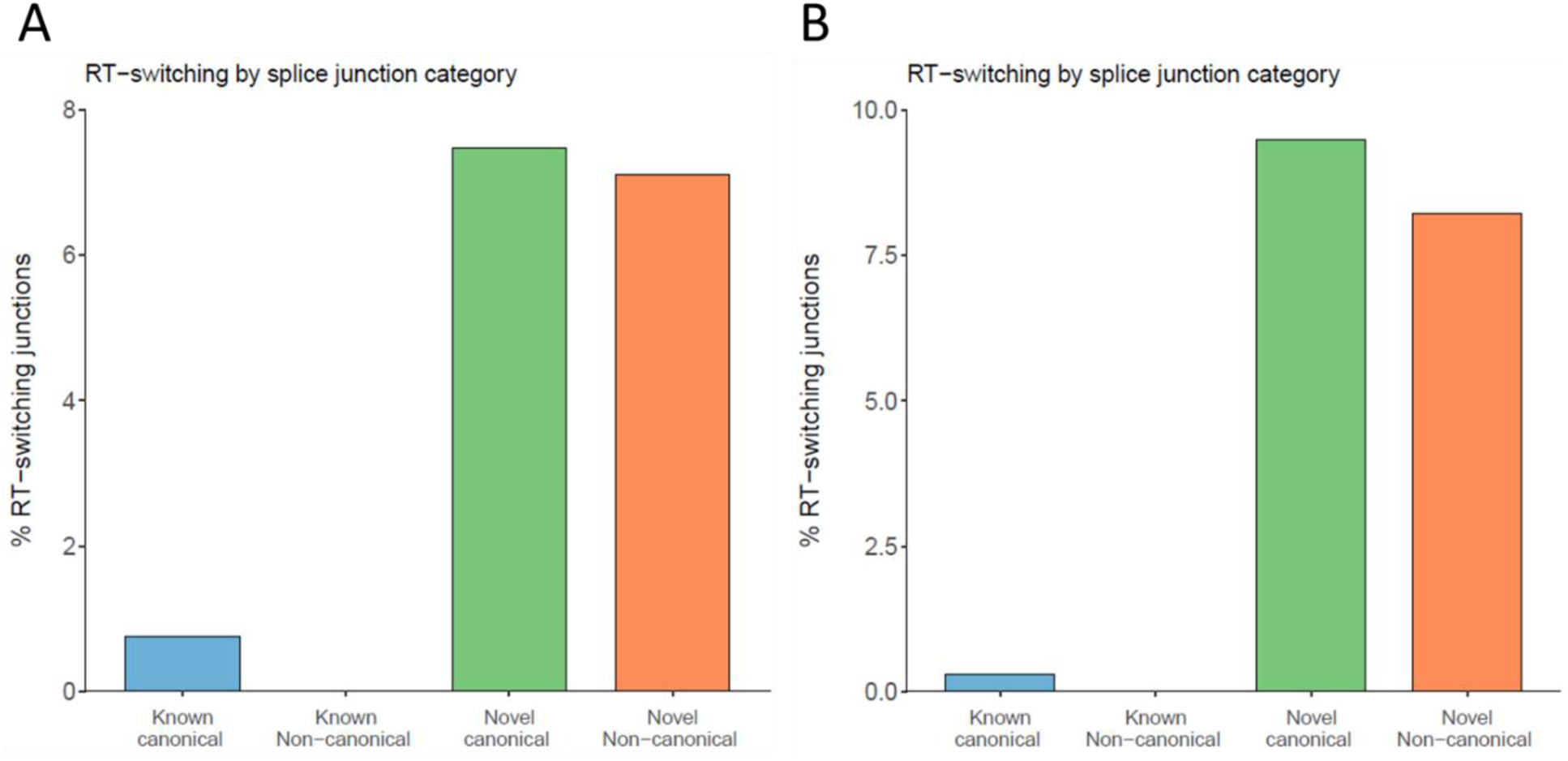
Identification of RT switching repeat sequence at splice junctions from different categories of splice junctions. A) Splice junction analysis of the human brain sample. B) Splice junction analysis of the human liver sample.

### PCR Amplification Bias

In many of the non-targeted RNA sequencing methods, full length cDNA is amplified after reverse transcription, typically to increase the DNA concentration for downstream library preparation like size selection and/or adaptor ligation. With LoopSeq Transcriptome kit, each unique cDNA molecule was assigned a molecular barcode during reverse transcription. The cDNA molecules were amplified for 20 PCR cycles, before the molecular barcodes were distributed across the length of the cDNA molecules and converted into sequencing short reads. By comparing the short read count binned to each unique cDNA molecules, the PCR amplification bias from full length cDNA amplification can be estimated. Figure 5 examines the short read count per unique molecule for the human brain sample, in comparison with three PCR bias simulations. As the PCR amplification bias increases, simulated by having a larger efficiency difference in the doubling rate across molecules within the PCR pool, the difference in the read count per molecule increases. With low amplification bias, there is less than one order of magnitude difference in the read count per molecule within the PCR pool. The difference increases to three orders of magnitude when the PCR amplification bias is severe. For the human brain sample, the PCR amplification bias closely resembles the medium amplification bias simulation for molecules with >200 reads but drifts towards high amplification bias for molecules with 100 reads or lower. One possible explanation for the larger than expected population of molecules with low read count is that an error in the barcode region is introduced either during cDNA amplification or short read sequencing, leading to a new barcode sequence being created and an appearance of a slow amplifying molecule. However, these molecules account for roughly 5% of the total reads, while the top 25% of the molecules account for roughly 48% of the total reads. The imbalance of molecule counts to read counts means the molecules with high copy numbers after cDNA amplification will be disproportionally sampled during short read and long read sequencing over molecules with lower copy numbers. Figure 5 underscores the importance of tagging the cDNA molecule with unique molecular barcodes prior to amplification. By tagging the cDNA molecules with unique molecular barcodes, one can differentiate between highly abundant, identical cDNA molecules and obtain accurate transcript abundance information, but can also differentiate unique cDNA molecules from PCR duplicates and assess the degree of amplification bias introduced during cDNA amplification.

**Figure 5.**
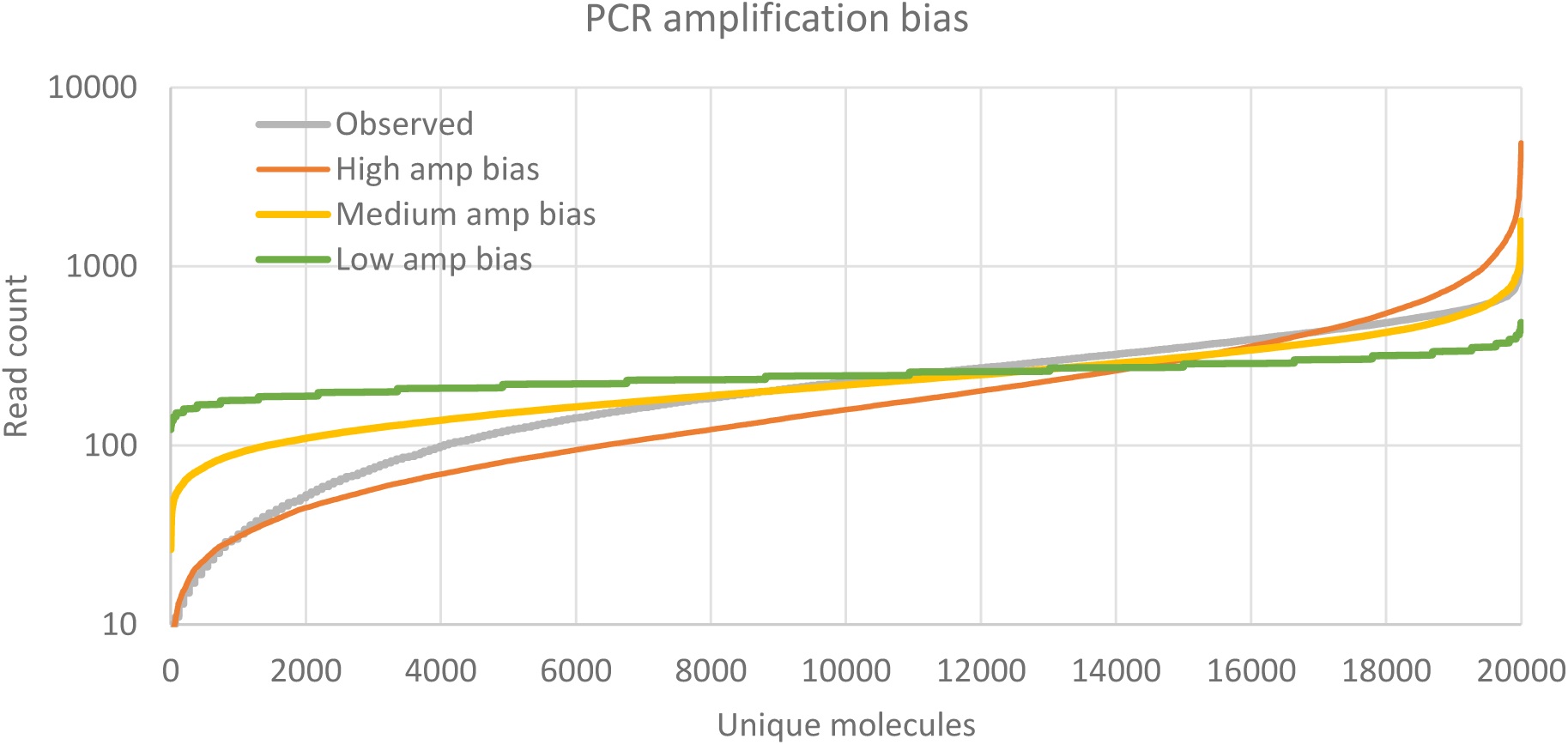
PCR amplification bias from cDNA amplification. The observed read count per unique molecule for the human brain sample is shown in grey, while the simulated PCR amplification biases are shown in orange (high amplification bias), yellow (medium amplification bias), and green (low amplification bias).

## Discussion

Full-length transcriptome sequencing using LoopSeq synthetic long read technology is an exciting approach to surveying the diverse transcripts and splice variants that exist across different biological samples. Here we have demonstrated full-length transcriptome sequencing of three human tissue extracted RNA using the LoopSeq Transcriptome kit. After library preparation and sequencing, synthetic long reads, or contigs, were reconstructed from short reads originating from single molecules as identified by the molecular barcodes. The *de novo* assembly of contigs relies on overlapping sequences within the pool of short reads that share the same molecular barcode. There are two implications to this. First, there exists a minimum short read coverage per transcript to provide enough coverage for full length assembly to be possible, if long read reconstruction and phasing is the intended goal. However, because there are many short reads covering each nucleotide position per transcript, the reconstructed contigs are the consensus sequences that correct for errors that introduced either during library amplification or during short read sequencing. When a single base pair variation or a short indel event is observed in the transcript, the higher the sequencing coverage per transcript there is, the higher the confidence one has that this is a true positive event. In addition, since each contig represents a unique cDNA molecule prepared during reverse transcription, one can also have high confidence in a specific splice variant when it is observed across multiple contigs.

There are many exciting applications of the LoopSeq Transcriptome kit and the LoopSeq synthetic long read technology. One application is in the area of target enrichment and sequencing only specific genes of interest. Sequencing whole transcriptome typically means devoting most of the sequencing reads to the highly abundant transcripts in the samples, due to the nature of single-molecule sequencing. Since the cDNA is already tagged with UMI, the target enrichment can be performed on the full-length cDNA molecules, enriching for transcript isoforms from specific genes of interest, before subjecting the cDNA molecules to LoopSeq library preparation. In addition to sequencing whole transcriptomes, the LoopSeq synthetic long read technology can be applied to sequencing cDNA molecules tagged with cell-specific barcodes from single-cell applications. Depending on the design of the single-cell barcode adaptor, the design of the LoopSeq barcode adaptor might need to be adjusted to facilitate downstream barcode distribution within the length of the cDNA molecules. Once the LoopSeq barcode is distributed randomly within the cDNA molecules and sequenced alongside internal regions of the cDNA molecules, the cDNA can then be reconstructed with their cell-specific barcodes. The junction between LoopSeq synthetic long read technology and RNAseq represent many exciting new opportunities and research angles to push the boundaries of human transcriptome sequencing.

## Materials and Methods

### Materials

Human brain total RNA was purchased from ThermoFisher Scientific, while human liver total RNA and human blood total RNA were purchased from Zyagen. LoopSeq Transcriptome kit was obtained from Loop Genomics. SPRIselect reagent was purchased from Beckman Coulter. RNA qualification and quantification using Bioanalyzer RNA Nano 6000 Kit was purchased from Agilent, and RNA quantification using Qubit RNA HS Assay Kit was purchased from ThermoFisher Scientific. Library quantification kit was purchased from Kapa Biosystems (Roche), and library qualification and quantification reagent using Bioanalyzer High Sensitivity Kit was purchased from Agilent. NextSeq 500/550 kit for short read sequencing was purchased from Illumina.

### Full-length cDNA synthesis, library preparation and sequencing

The full-length cDNA synthesis, cDNA amplification, as well as short read sequencing library preparation were completed using the LoopSeq Transcriptome 3x 8-plex kit. Briefly, 10 ng of total RNA from human brain, liver, and blood total RNA were reverse transcribed using polythymine oligoes and tagged with sample barcodes and molecular barcodes (UMI). After reverse transcription, samples undergo an enzymatic cleanup as well as a SPRIselect cleanup. The purified cDNA was amplified and pooled into a single reaction for downstream processing. Different amounts of barcoded cDNA were used as input in the cDNA amplification reactions, depending on the number of unique tagged cDNA intended for transcriptome sequencing. In this experiment, approximately 5,000 – 33,000 uniquely tagged cDNA molecules were amplified, which is equivalent to <1% of the total cDNA synthesized. After cDNA amplification, the molecular barcodes were then distributed across the length of the cDNA and underwent another enzymatic cleanup and SPRIselect cleanup. The cleaned product was fragmented, ligated with Illumina adaptors, and amplified with an Illumina sample index. The final library was purified using SPRIselect reagent, and QC’d using Agilent Bioanalyzer High Sensitivity assay as well as Kapa library quantification kit. Short read sequencing was obtained on an Illumina NextSeq machine, with a sequencing depth of 3M to 7M PE150 reads per sample.

### Data analysis

The fastq output files from sequencing was used as the input into the Loop Genomics data analysis pipeline for sample demultiplexing and long read reconstruction. Briefly, the fastq files were verified and demultiplexed using the sample barcodes. The short reads from each sample were then split into chunks using molecular barcodes, before being reconstructed into single-molecule long reads using *de novo* assembly algorithm. The output long reads, or contigs, were reference mapped to the human RefSeq RNA transcript database for sequence identification and for generating sample prep statistics. SQANTI analysis was conducted on the long read contig outputs from the Loop Genomics pipeline using reference human genome GRCh38 and ENSEMBL genome annotation file for sequence alignment.

### PCR amplification bias simulation

To simulate the effect of different PCR amplification bias on sequencing read sampling, the theoretical copy numbers of 20,000 unique molecules with different doubling rates were calculated. Three doubling rates were examined: a random doubling rate between 1 and 2 at every PCR cycle for the high amplification bias, between 1.4 and 2 for the medium amplification bias, and between 1.8 and 2 for the low amplification bias sample. 5M random reads were selected from the pool of molecule copies generated from 20,000 unique molecules after 20 PCR cycles and plotted against the observed copy number per unique molecule from the human brain RNA sample (medium sequencing depth).

